# A western representative of an eastern clade: phylogeographic history of the gypsophilous plant *Nepeta hispanica*

**DOI:** 10.1101/2022.01.05.475097

**Authors:** Ignacio Ramos-Gutiérrez, Juan Carlos Moreno-Saiz, Mario Fernández-Mazuecos

## Abstract

The preference of certain plant species for gypsum soils leads to disjunct population structures that are thought to generate island-like dynamics potentially influencing biogeographic patterns at multiple evolutionary scales. Here, we study the evolutionary and biogeographic history of *Nepeta hispanica*, a western Mediterranean plant associated with gypsum soils and displaying a patchy distribution with populations very distant from each other. Three approaches were used: (a) interspecific phylogenetic analyses based on nuclear DNA sequences of the ITS region to unveil the relationships and times of divergence between *N. hispanica* and its closest relatives; ((b)phylogeographic analyses using plastid DNA regions *trn*S-*trn*G and *psb*J-*pet*A to evaluate the degree of genetic isolation between populations of *N. hispanica*, their relationships and their genetic diversity; and (c) ecological niche modelling to evaluate historical distributional changes. Results reveal that *N. hispanica* belongs to an eastern Mediterranean and Asian clade diversified in arid environments since the Miocene-Pliocene. This species represents the only extant lineage of this clade that colonized the western Mediterranean, probably through the northern Mediterranean coast (southern Europe). Present Iberian populations display a high plastid genetic diversity and, even if geographically distant from each other, they are highly connected according to the distribution of plastid haplotypes and lineages. This can be explained by a scenario involving a complex history of back-and-forth colonisation events, facilitated by a relative stability of suitable conditions for the species across the Iberian Peninsula throughout the Quaternary.

## Introduction

Studying the eco-evolutionary history of plant species helps us to understand how they have reached their present distribution and population structure. A key factor to consider is soil composition, which often determines the capacity of species to thrive in a certain area. A paradigmatic example of this effect is the flora linked to gypsum soils, which exert an extraordinary selective pressure on plant species, leading to the evolution of floras highly adapted to them (Kruckeberg, 1986). These floras frequently display peculiar biogeographic patterns due to the patchy distribution of gypsum soils at different scales, and this is particularly true for the Mediterranean region. (1) At a regional scale, several gypsum-associated plant lineages show a wide disjunction, in which some representatives are distributed in the western Mediterranean, while their closest relatives are found in the eastern Mediterranean. This distribution pattern has been historically interpreted as the result of vicariance following the Messinian Salinity Crisis during the late Miocene, when the Mediterranean sea level drastically dropped, providing an ecological corridor that allowed plants associated with salty environments to travel westwards across the Mediterranean basin (Bocquet et al., 1978). (2) At a local scale, gypsum soils are found in continental areas, and usually form island-like patches surrounded by other substrates. This creates a unique population structure in gypsum-associated plants, in which patterns of gene flow and geographic isolation may be similar to those of island plants (Moore et al., 2004).

*Nepeta* L. is a genus of the mint family (Lamiaceae) which is distributed throughout temperate Eurasia and northern Africa and includes several gypsum-associated species. With nearly 300 species in total, most of them perennial herbs, it is one of the most diverse genera of the family (Ubera & Valdés, 1983; Jamzad et al., 2003b). The highest species diversity is found in south-western Asia (Jamzad et al., 2003a). Within *Nepeta*, the section *Oxynepeta* is characterized by presenting dioecy, and comprises five currently accepted species (Budantsev, 1993; POWO, 2021). Of these, the type species is found in eastern Europe and south-western Asia (*Nepeta ucranica* L.), three more species are found in south-western Asia (*N. congesta* Fisch. & Mey., *N. heliotropifolia* Lam., and *N. stricta* (Banks & Sol.) Hedge & Lamond), and only one species is found in the western Mediterranean Region, including the Iberian Peninsula and north-western Africa (*N. hispanica* Boiss. & Reut.). A detailed phylogenetic study of this section has not yet been carried out, and only two species (*N. heliotropifolia* and *N. congesta*) were included in the most comprehensive study of the genus undertaken so far (see Jamzad et al., 2003b). Given the limited availability of phylogenetic data and the vast geographical distance between *N. hispanica* (native to the western Mediterranean) and the rest of *Oxynepeta* species (with Irano-Turanian and East Mediterranean distributions) there remains the possibility that dioecy (the diagnostic character of sect. *Oxynepeta*) evolved more than once during the diversification of the genus (Gleiser & Verdú, 2005; Schaefer & Renner, 2010), thus meaning that the sectional classification does not reflect the evolutionary history of the group.

*Nepeta hispanica* Boiss. & Reut. is a western Mediterranean endemic species that shows the peculiar biogeographic patterns described above for gypsum-associated species at both geographic scales: an eastern-western Mediterranean disjunction with its putative closest relatives, and a patchy distribution of populations. On the one hand, based on morphological traits, some authors have suggested a very close relationship between *N. hispanica* and *N. ucranica* (Aedo, 2010), or even considered the former a subspecies of the latter (*Nepeta ucranica* subsp. *hispanica* (Boiss. & Reut.) Bellot, Casaseca & Ron). On the other hand, *N. hispanica* occurs in patchy populations that are also distant from one another as a result of its preference for gypsum soils (de la Cruz et al., 2011). This scattered distribution, with populations separated by hundreds of kilometres, extends from the inner Iberian Peninsula to northern Morocco. Some taxonomists (Ubera & Valdés, 1983) separated central-northern Iberian populations as a different species (*N. beltranii* Pau, currently considered a synonym of *N. hispanica*; Aedo, 2010), while Moroccan populations are recognised as *N. hispanica* subsp. *statice* Maire (Fennane et al., 1998). Nevertheless, the phylogenetic relationships of *N. hispanica* within the genus and the genetic variability and relationships between populations remain unstudied. Apart from its biogeographic and evolutionary interest, advancing the knowledge on this species is important because it is a threatened and protected plant in several territories (de la Cruz et al., 2011).

Here we used three approaches to shed light on the evolutionary and biogeographic history of *N. hispanica*. First, we conducted phylogenetic analyses to test the monophyly and estimate divergence times of *Nepeta* sect. *Oxynepeta* by combining previously published and newly generated ITS sequences (including those of *N. hispanica*). Second, we performed a phylogeographic analysis of *N. hispanica* based on plastid DNA (ptDNA) sequences to evaluate the degree of genetic isolation between populations, the relationships between them and their genetic diversity. And third, we modelled the present environmental niche of the species and projected it to past periods to help interpret phylogeographic patterns and historical distributional changes. We tested the following hypotheses: (1) sect. *Oxynepeta* is a monophyletic group, and therefore *N. hispanica* is closely related to other dioecious species of the genus; and (2) long-term geographic isolation, resulting from a discontinuity in historically suitable areas, has led to a strong genetic isolation between populations of *Nepeta hispanica*.

## Methods

### Sample collection

For the phylogenetic analysis, we sampled herbarium specimens of the three species of *Nepeta* sect. *Oxynepeta* with no DNA sequences available from previous studies, including two geographically distant specimens of *N. hispanica*, one of *N. ucranica* and one of *N. stricta* (Table S3). All of them were provided by the Royal Botanical Garden of Madrid Herbarium (MA). For each species, the most recently collected specimens were used to ensure DNA quality.

For the phylogeographic analysis, we prospected all 14 known Iberian localities of *Nepeta hispanica*. Since the species appears to have become extinct in some of them, we ended up collecting leaf tissue of a total of 111 individuals from 10 populations (12 individuals per population, except one in which only three individuals were found). Whenever possible, similar numbers of male and female individuals were collected. Samples were kept in paper envelopes and dried in silica gel. In populations present in regions where the species was not legally protected, herbarium specimens were prepared and deposited at the Royal Botanical Garden of Madrid (MA) and Universidad Autónoma de Madrid (MAUAM) herbaria. Moroccan populations of *N. hispanica* subsp. *statice* could not be visited, but we obtained one sample provided the Geneva Herbarium (G) (No. J7635), now deposited at the MA Herbarium.

### DNA sequencing

Genomic DNA was extracted from all samples using the *DNeasy Plant Mini Kit* (Qiagen, USA) following the manufacturer’s protocol, except that incubation time varied depending on the sample (up to 1.5 h for herbarium samples).

For the phylogenetic analysis, we selected the internal transcribed spacer (ITS) region, which was used in the most extensive phylogenetic analysis of *Nepeta* published to date (Jamzad *et al*., 2003b). The ITS region was amplified by PCR and sequenced for the *N. hispanica, N. ucranica* and *N. stricta* samples using the external primers 17SE-26SE and the internal primers P1A-P4 (Sun et al., 1994; Jamzad et al., 2003b). However, a good-quality sequence of *N. stricta* could not be obtained, possibly due to fungus contamination.

For the phylogeographic analysis of *Nepeta hispanica*, we first performed a pilot study to select consistently amplified and variable ptDNA regions. Seven regions of the plastid genome, considered potentially informative for angiosperms (Shaw *et al*., 2007), were sequenced for one individual per population. As a result, we selected the two regions (*trnS-trnG* and *psbJ-petA*) with the highest numbers of SNPs (single nucleotide polymorphisms). Then these two ptDNA regions were amplified by PCR for ten individuals per population of *N. hispanica*, and for the *N. ucranica* sample to be used as outgroup in the haplotype analysis. Standard primers were used, in combination with newly designed internal primers for samples in which amplification proved difficult (Table S2).

In all cases, PCR was conducted using BIOTAQ DNA polymerase (Bioline, USA). PCR conditions consisted of an initial denaturation step at 94°C for 1 min and 30 cycles of amplification, each consisting of denaturation at 94° for 1 min, annealing at 50-52°C for 1 min, and extension at 72°C for 1 min, followed by a final extension step at 72°C for 10 min. PCR products were checked by electrophoresis in 1% agarose and submitted to Macrogen Inc. (Madrid, Spain) for Sanger sequencing. Sequences were assembled and edited using Geneious 11.1.13 (Kearse et al., 2012). All sequence variation among individuals was double-checked on electropherograms to prevent mistakes.

### Phylogenetic analysis and dating

For phylogenetic analyses, the three newly generated ITS sequences (two belonging to *N. hispanica* and one to *N. ucranica*) were combined with those included in the analysis of Jamzad et al. (2003b), obtained from the GenBank database (Benson *et al*., 2005). These included sequences of two additional species of *Nepeta* sect. *Oxynepeta* (*N. heliotropifolia* and *N. congesta*), 31 other species of *Nepeta* and 5 closely related genera to be used as the outgroup. A total of 36 individuals of 35 taxonomically representative species of the genus *Nepeta* were included. All ITS sequences were aligned using the MAFFT algorithm (Katoh et al., 2002), one of the quickest and more precise among those currently available. The sequence matrix was analysed using the jModelTest software (Posada, 2008) to estimate the best-fitting nucleotide substitution model (*GTR+I+G*). A phylogenetic tree based on Bayesian inference was obtained using MrBayes 3.2 (Huelsenbeck & Ronquist, 2001). Two parallel runs of 10 million generations each were performed, and sampled every 1000 generations. A majority-rule consensus tree was calculated with a burn-in phase of 10%. Mixing and convergence were checked using Tracer 1.7 Rambaut et al., 2018).

To obtain a time-calibrated phylogeny, we analysed the ITS matrix (including a single specimen per species) using BEAST2 (Bouckaert et al., 2014). A secondary calibration based on previous information from Hermant et al. (2012), Drew & Sytsma (2012) and Deng *et al*. (2015) was implemented. This information was obtained from the TimeTree database (www.timetree.org; Kumar *et al*., 2017), in which the separation of the genera *Nepeta* and *Agastache* (included in the outgroup) was estimated to have happened between 25.7 and 31.3 million years ago. This interval was used as a uniform prior for the root of the tree. A birth-death process and an uncorrelated lognormal relaxed clock were implemented. Four chains of 10 million generations were run and sampled every 1000 generations. Tracer was used to confirm mixing and convergence, and determine an appropriate burn-in fraction. After the combination of chains using LogCombiner (with a burn-in phase of 10%), a maximum clade credibility (MCC) tree was built in TreeAnnotator (both programs are part of the BEAST2 package) and visualized using FigTree 1.4 (Rambaut, 2014).

### Phylogeographic analyses

Sequences of the two ptDNA regions (*trnS-trnG* and *psbJ-petA*) for the 94 individuals of *N. hispanica* and the outgroup individual of *N. ucranica* were separately aligned using MAFFT. For phylogeographic analyses, both ptDNA regions were concatenated. To estimate relationships among haplotypes, a haplotype network was inferred using the statistical parsimony method (Templeton, 1992) implemented in the TCS 1.21 software (Clement et al., 2000), with the *gaps* = *missing* option and a connection limit of 95%. The distribution of haplotypes across populations was mapped using QGIS 3.4.2 (QGIS Development Team, 2018).

Additionally, a discrete phylogeographic analysis (hereafter DPA; Lemey et al., 2009) was carried out to shed light on migration patterns among biogeographic areas. This analysis was carried out in BEAST 1.10 (Suchard et al., 2018). Five discrete areas were defined, including the four Iberian river basins in which *Nepeta hispanica* is found (namely Ebro, Duero, Tagus and Guadalquivir river basins) and northern Africa. *N. ucranica* was used as outgroup with an undetermined area (because we were only interested in migration within *N. hispanica*). An asymmetric substitution model was implemented for the area partition, while substitution model for the DNA partitions followed jModelTest results. A strict clock was used for the area partition and an uncorrelated relaxed clock with a lognormal distribution for the DNA partitions, with a constant size coalescent tree prior. The root age was calibrated using a uniform prior distribution between 2.11 and 7.7 Mya based on the divergence time estimated by the dating analysis of ITS sequences (see Results). Two MCMC chains of 10 million generations sampled every 1000 generations were run, and they were combined in LogCombiner with a burn-in phase of 10%. Mixing and convergence of chains was confirmed using Tracer 1.7. A MCC tree was built in TreeAnnotator considering common ancestor heights. To evaluate the statistical support of migration routes, Bayesian stochastic search variable selection (BSSVS) was implemented, and Bayes Factors (BFs) were calculated in SPREAD3 (Bielejec et al., 2016).

### Ecological niche modelling

For ecological niche modelling, a maximum entropy algorithm was used (Phillips *et al*., 2006) because it is considered to work well with presence-only data (Elith *et al*., 2011). To prevent geographic bias, the analysis was performed only for the Iberian Peninsula, where the distribution range of *N. hispanica* is well known. Northern Africa was not included because of the limited available information on the species distribution in this region. Occurrence points included those obtained during our own fieldwork and those obtained from the biodiversity database GBIF (www.gbif.org, 2018). However, we only used high-accuracy occurrence data postdating 1901 (at least 4 decimal places in their coordinates, an error corresponding to less than 10 m at that latitude). Oversampling issues (redundant data in more accessible areas) were corrected by the random elimination of points that were less than 0.05° apart from each other using a custom R script.

Bioclimatic, edaphic and lithological variables at 30 arc-second resolution were included in the model. Bioclimatic data were obtained from the CHELSA database (https://chelsa-climate.org; Karger et al., 2017), whereas edaphic data were taken from the SoilGrids database (https://soilgrids.org/; Hengl et al., 2017). To avoid collinearity issues in these sets of variables, only non-redundant ones were selected. To that end, a two-step procedure was implemented using the R packages “raster” (Hijmans, 2018) and “HH” (Heiberger, 2020) for bioclimatic and edaphic variables separately. First, we selected variables with a correlation below 70% in the study area (Dormann et al., 2013). Then, we excluded additional variables with a high variance inflation factor (VIF) until all remaining variables had VIF < 10. Selected variables are shown in Table 1. Additionally, a lithological categorical variable, mapping nine lithological classes potentially influencing plant distributions (Fernández-Mazuecos & Glover, in prep.) was included because *N. hispanica* is known to be associated with gypsum soils. This lithological variable is based on the Lithostratigraphic Map of Spain (del Pozo Gómez, 2009). Maximum entropy modelling was implemented in MaxEnt 3.4.1 (Phillips *et al*., 2006). The model was generated by averaging 100 *subsample* replicates, each one being assigned 90% of point data for modelling and 10% for validation.

**Table 1.** Bioclimatic, edaphic and lithological variables included in the ecological niche model of *Nepeta hispanica*, sorted by their percent contributions to the model. Permutation importance and effect direction are also shown for every variable.

| Variable | Variable description | Percent contribution | Permutation importance | Effect direction |
| --- | --- | --- | --- | --- |
| bio16 | Precipitation of wettest quarter | 35.1 | 82.2 | Sigmoid-negative |
| eda09 | pH index measured in water solution | 22.4 | 0.0 | Neutral |
| bio04 | Temperature seasonality | 19.0 | 5.9 | Linear-positive |
| iberlit | Lithology | 19.0 | 7.4 | Preference for categories “gypsic” and “fine-grained clastic”. |
| bio11 | Mean temperature of coldest quarter | 2.7 | 1.6 | Hump-shape |
| eda06 | Cation exchange capacity of soil | 0.6 | 1.3 | Sigmoid-positive |
| eda03 | Weight percentage of clay particles (<2·10<sup>-4</sup> mm) | 0.4 | 0.5 | Linear-negative |
| bio17 | Precipitation of driest quarter | 0.3 | 0.1 | Almost neutral-negative |
| eda02 | Volumetric percentage of coarse fragments (>2 mm) | 0.3 | 0.7 | Almost neutral-positive |
| bio08 | Mean temperature of wettest quarter | 0.1 | 0.3 | Neutral |
| eda07 | Soil organic carbon content | 0.1 | 0.1 | Sigmoid-negative |
| eda05 | Weight percentage of sand particles (0.05–2 mm) | 0.0 | 0.0 | Neutral |
| bio09 | Mean temperature of driest quarter | 0.0 | 0.0 | Neutral |

To estimate the areas with environmentally favourable conditions for *N. hispanica* in the past, the present model was projected to nine past time slices. Layers of bioclimatic variables for these periods at 2.5 arc-minute resolution were taken from the PaleoClim database (http://www.paleoclim.org/; see Table S4; Brown et al., 2018; Fordham et al., 2017; Karger et al., 2017; Otto-Bliesner et al., 2006). For these projections, only climatic conditions were considered to vary, while edaphic and lithological conditions were assumed to have stayed constant throughout these geologically recent periods.

## Results

### Phylogenetic analysis and dating

The phylogeny of 35 species of *Nepeta* obtained using the ITS region is shown in Figure 1. The four species of section *Oxynepeta* included in the analysis were recovered as a monophyletic group with a high posterior probability (PP=0.96). Within this clade, *N. ucranica* was recovered as the earliest-diverging species, with the remaining three species (*N. hispanica, N. congesta* and *N. heliotropifolia*) forming a moderately supported clade (PP=0.86).

**Figure 1.**
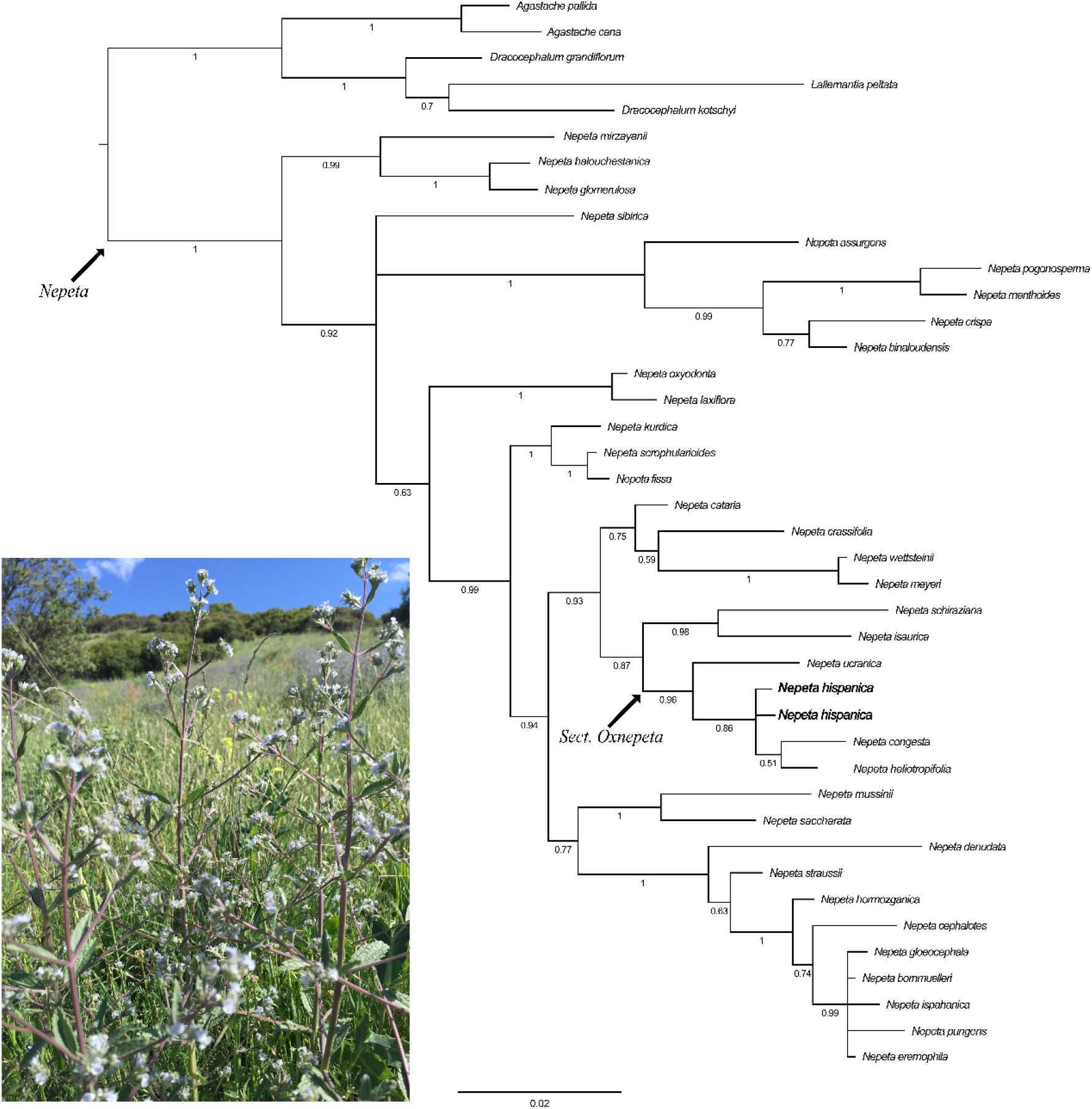
Phylogenetic position of *Nepeta hispanica*. The majority-rule consensus tree obtained in the Bayesian phylogenetic analysis of ITS sequences of 35 species of the genus *Nepeta* and five outgroup species is shown. Numbers below branches indicate posterior probabilities. The clades representing the genus *Nepeta* and section *Oxynepeta* are indicated with an arrow.

The dated ITS phylogeny of *Nepeta* sect. *Oxynepeta* and close relatives is shown in Figure 2. The sect. *Oxynepeta* clade was estimated to have split from its sister group at some point between the late Miocene and the early Pliocene (highest posterior density, HPD: 3.6-9.8 Ma; PP=0.83). The most recent common ancestor (MRCA) of the *Oxynepeta* species had an estimated age between the late Miocene and the early Pleistocene (HPD: 2.11-7.7 Ma; PP=0.95), while the common ancestor of *N. hispanica, N. congesta* and *N. heliotropifolia* had an estimated age between the late Pliocene and the Pleistocene (HPD: 0.59-3.98 Ma; PP=0.96).

**Figure 2.**
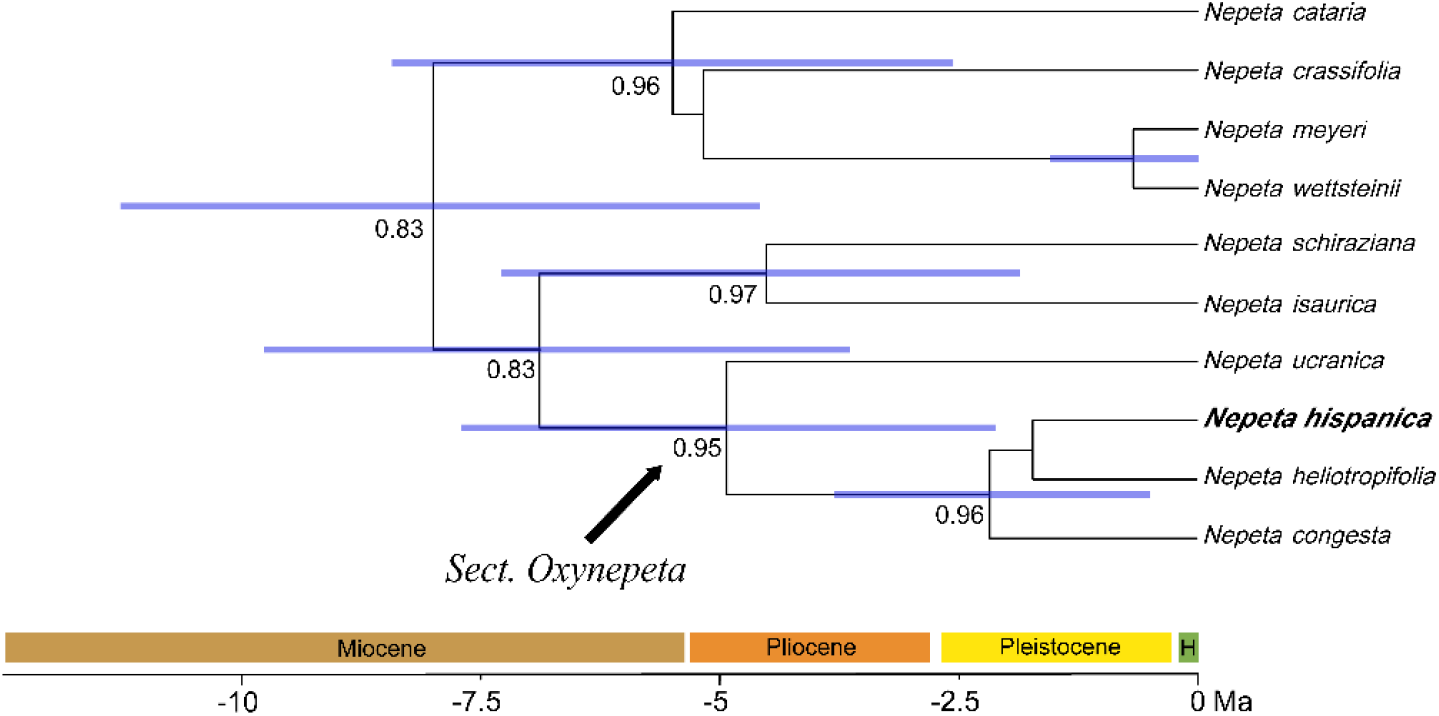
Dated phylogeny of *Nepeta* sect. *Oxynepeta* and close relatives. For clarity, other lineages included in the analysis are not shown. Numbers below branches are Bayesian posterior probabilities. Bars indicate 95% highest posterior density intervals for divergence times.

### Phylogeographic analyses

The geographic distribution and relationships of ptDNA haplotypes are shown in Figure 3. A high intraspecific haplotype diversity (16 extant haplotypes; 22 missing haplotypes) and a high number of nucleotide substitutions were detected. In the haplotype network (Figure 3b), haplotype A was interpreted as ancestral given that it is linked to the outgroup species (*N. ucranica*). It is also the most frequent haplotype, widely distributed across central and eastern Spain (five populations), and it shows the highest number of connections. From this ancestral haplotype, eight derived lineages radiate, each of them including between one and four haplotypes separated from the ancestral haplotype by one to six substitutions. The B1-B3 lineage formed a loop with the ancestral haplotype.

**Figure 3.**
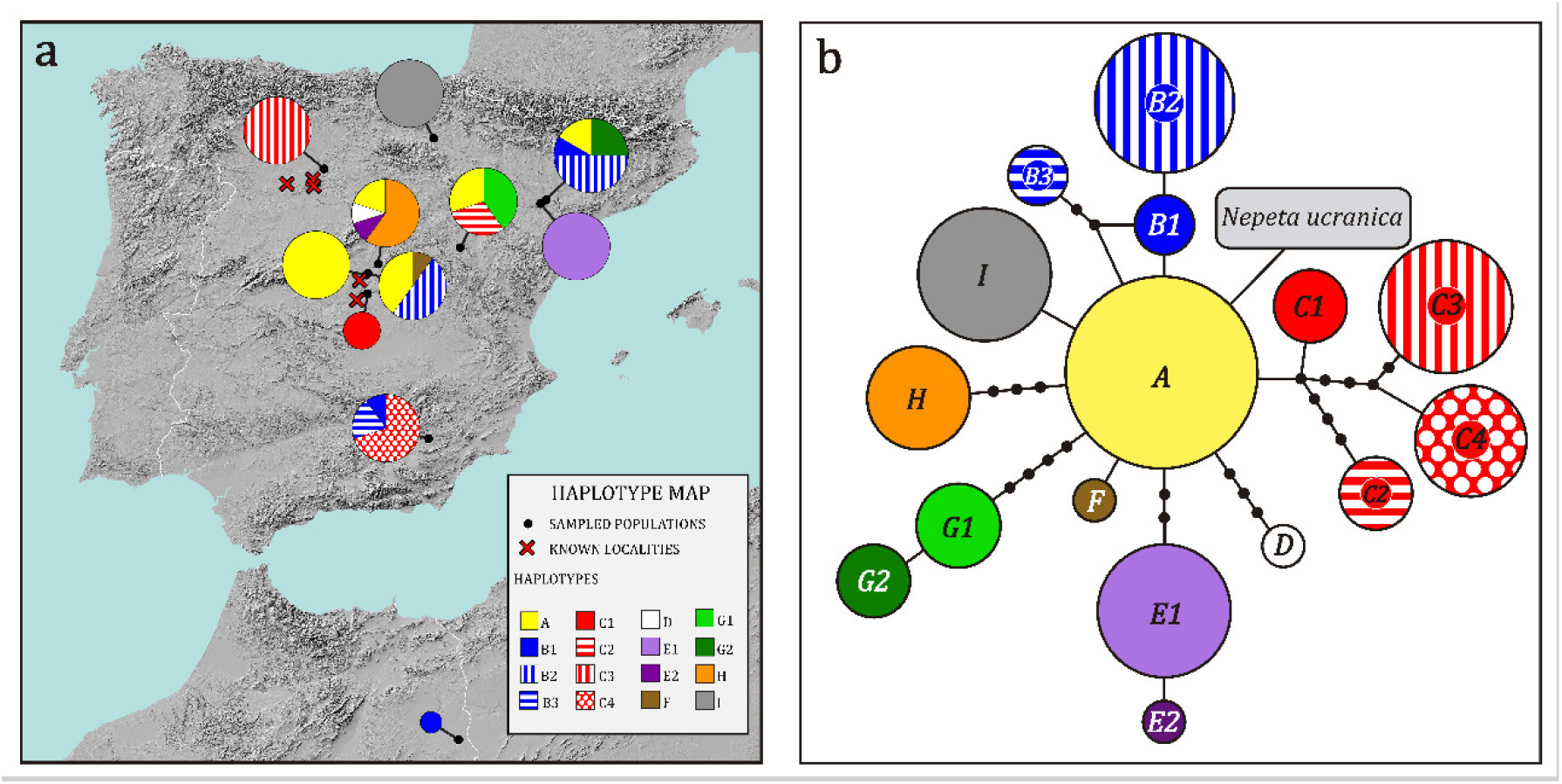
Phylogeographic analysis of *Nepeta hispanica* based on plastid DNA regions *trnS-trnG* and *psbJ-petA*. Each colour represents a haplotype. (a) Geographic distribution of haplotypes across the Iberian Peninsula and northern Africa. Pie charts indicate the frequencies of haplotypes in populations. Pie chart sizes are proportional to the number of sampled individuals per population. (b) Haplotype network. Circle sizes are proportional to haplotype frequencies. Each line represents a nucleotide substitution, and black dots represent missing haplotypes (extinct or not sampled).

The geographic distribution of haplotypes (Figure 3a) shows that the same haplotype, or phylogenetically close ones, can be found in very distant populations. For example, the derived haplotype B2 is found in two populations, one of them in the Tagus basin (central Spain) and the other in the Ebro basin (northeastern Spain). More strikingly, haplotype B1 is found in that same northeastern locality as well as in the Guadalquivir region (southeastern Spain) and the single Moroccan sample. Furthermore, very distant haplotypes can be found in the same or geographically close populations. For example, the population in Guadalquivir basin contains the distant haplotypes C4 and B1/B3, and a population in Duero’s one (northeastern Spain) contains the ancestral haplotype A together with the distant derived haplotypes G2 and B1/B2. Regarding diversity patterns, the highest numbers of haplotypes were found in central, northeastern and southeastern Spanish populations (five populations with 3-4 haplotypes per population, frequently from distant lineages). Five Spanish populations had no haplotype diversity (a single haplotype per population), two of them northern, two central and one northeastern.

The results of the DPA are shown on Figure 4. The most probable location of the most recent common ancestor of all plastid lineages of *Nepeta hispanica* was the Ebro river basin in northeastern Iberia (PP=0.59), followed by the Tagus basin in central Iberia (PP=0.36). According to Bayes factors, the best-supported migration route was that from the Ebro basin to the Tagus basin (BF=25.1), followed by the route from the Ebro basin to the Guadalquivir basin (BF=10.2). Six other migration routes received positive support (BF>3; Kass & Raftery, 1995), connecting adjacent and distant areas.

**Figure 4.**
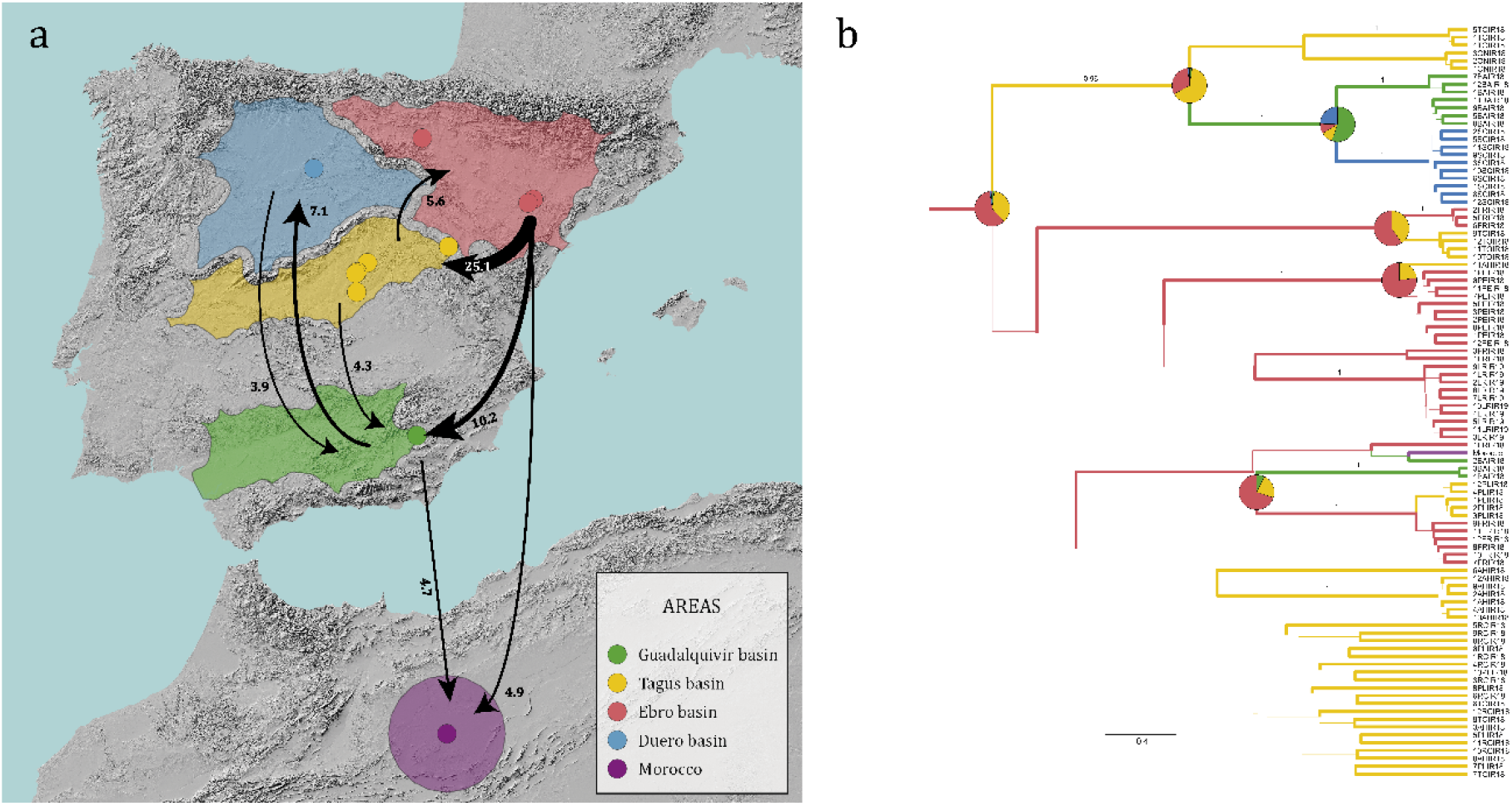
Discrete phylogeographic analysis (DPA) of *Nepeta hispanica* in the Iberian Peninsula and North Africa. (a) Map of sampled populations in the five geographic areas (five river basins in the Iberian Peninsula, and northern Africa. Arrows represent routes of migration between areas with Bayes factor (BF) values above 3, with numbers and width indicating BF values. (b) DPA consensus tree, in which branch widths represent posterior probabilities, and colours indicate areas as in (a). Pie charts show probabilities of ancestral areas for nodes with posterior probability above 0.9. The outgroup is excluded for clarity.

### Ecological niche modelling

Figure 5 shows the result of the maximum entropy distribution modelling, both for the present and for past time slices. Percent contributions and permutation importance of variables are shown in Table 1. According to these, the most relevant variables included climatic (precipitation of wettest quarter, temperature seasonality), edaphic (soil pH) and lithological variables. Present suitable areas for *N. hispanica* were estimated throughout the eastern half of the Iberian Peninsula, with the highest suitability values in areas with continental climate and gypsic or fine-grained clastic lithologies (gypsum, gypsiferous marl, lutite, silt, clay). Projections to past time slices showed a relatively constant distribution range, with some waxing and waning. In particular, the greatest range reduction was obtained for the Last Glacial Maximum (LGM), when estimated suitable areas were confined to the eastern continental areas of the Iberian Peninsula.

**Figure 5.**
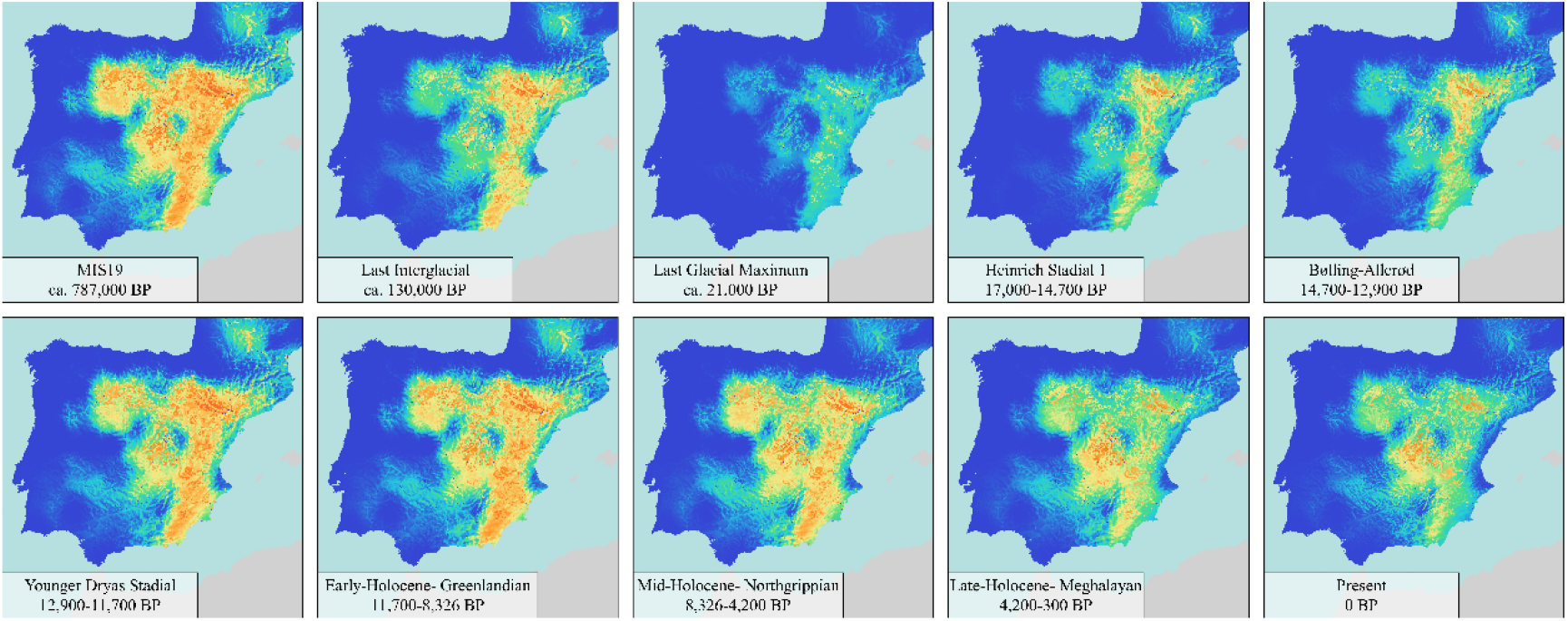
Ecological niche modelling of *Nepeta hispanica* in the Iberian Peninsula. Environmental suitability values for the present and nine past time slices according to maximum entropy modelling are represented, with colder colours indicating low suitability, and warmer colours indicating higher suitability.

## Discussion

This study provides a two-scale picture of the evolutionary and biogeographic history of the gypsum-associated Mediterranean species *Nepeta hispanica*. On the one hand, phylogenetic results confirm a close relationship between the western Mediterranean *N. hispanica* and the other geographically distant species of *Nepeta* sect. *Oxynepeta*, distributed in the eastern Mediterranean and Irano-Turanian regions. On the other hand, phylogeographic results based on plastid DNA revealed a remarkable lack of genetic isolation between the distant island-like patches making up the distribution range of *N. hispanica*.

### A western Mediterranean representative of a predominantly eastern clade

Phylogenetic analyses are congruent with the classical morphological criterion used to define *Nepeta* sect. *Oxynepeta* (Budantsev, 1993). Indeed, results support that dioecy, the diagnostic character of the section, has evolved only once in the genus *Nepeta*. As a result, the four sampled species of sect. *Oxynepeta* (out of a total of five currently accepted species) form a monophyletic group in our ITS phylogeny. The common ancestor of this group is estimated to have split from its sister lineage around the late Miocene to early Pliocene, possibly in the eastern Mediterranean or Irano-Turanian region, based on the distribution ranges of species in the sister lineage (*N. isaurica, N. schiraziana*) and of the earliest-diverging species of sect. *Oxynepeta* (*N. ucranica*). It is also estimated that, at some point in the last 4 million years, a lineage of sect. *Oxynepeta* migrated to the western Mediterranean. This lineage eventually became isolated and gave rise to a distinct species, *N. hispanica*. Therefore, *Nepeta hispanica* stands as the only western-Mediterranean representative of this otherwise eastern clade. This longitudinal Mediterranean disjunction has been widely discussed for different species. Although it has long been hypothesized that migrations from the eastern Mediterranean and the Irano-Turanian region to the western Mediterranean occurred mainly in the Messinian Age (7.2-5.3 Ma), coinciding with the Messinian Salinity Crisis (Bocquet et al., 1978), the divergence of *N. hispanica* appears to have happened later, and therefore may be the result of long-distance dispersal between environmentally similar eastern and western habitats. It should be noted that our phylogeny (based on that of Jamzad et al., 2003) has been obtained using a single DNA region and a limited sample of *Nepeta* species, and divergence time estimates have wide confidence intervals. While divergence times should be taken with caution, we consider that the close relationship between *N. hispanica* and the eastern species of sect. *Oxynepeta* is well supported based on the congruence of ITS sequences and morphology. A more deeply sampled time-calibrated phylogeny of *Nepeta* based on multi-locus data combined with explicit biogeographic analyses will be needed to get deeper insights into the diversification history of Mediterranean and Irano-Turanian *Nepeta* lineages.

### Lack of phylogeographic structure despite an island-like range

We hypothesize that the migration and arrival of the ancestor of *N. hispanica* in the western Mediterranean from the east (eastern Mediterranean and Irano-Turanian regions) happened along the northern Mediterranean coast (southern Europe). Indeed, the most likely ancestral areas for *N. hispanica* populations are found in central and northern Iberia, and the haplotype of the only sample from northern Africa (Morocco) is derived from an ancestral Iberian haplotype and is shared with several Iberian populations. Colonisation of northern Africa was likely the result of a relatively recent event of long-distance dispersal from the Iberian Peninsula (Terrab et al., 2008 Blanco-Sánchez et al., 2021). However, additional northern African samples would be necessary to confirm this hypothesis and evaluate the evolutionary history of the species in this region. Because of this lack of information about northern African populations, here we have focused on disentangling the phylogeographic structure of the better-known Iberian populations.

Since the establishment of *N. hispanica* in the Iberian Peninsula, estimated to have happened during the late Pliocene or Pleistocene (not before 4 Ma), a history of sequential colonisation of areas started, with substantial migration between areas in multiple directions, as revealed by ptDNA lineages. Throughout Iberian populations, high variability and low phylogeographic structure of ptDNA sequences is observed at the global and population scales, which is congruent with a history of genetic connection rather than isolation. Species distribution modelling revealed a relatively wide extension and temporal continuity of available habitats with continental climates and suitable soils throughout the late Quaternary. This appears to have allowed the preservation of high genetic variation and facilitated long-term genetic exchange between populations. The only period in which this continuity was partially interrupted is the LGM (approximately 21,000 years ago, cf. Figure 5). However, suitable areas remained in the inner-eastern areas of the Iberian Peninsula, where populations thrive nowadays. These areas therefore may have acted as genetic repositories from where the remaining populations were recolonised. This hypothesis is in line with DPA results, in which Ebro basin populations are estimated as the ancestral ones, from where other populations were colonised in a series of back-and-forth events. A general north-to-south pattern has been described for other Iberan gypsophytes (Salmerón-Sánchez et al., 2017), and may be explained by a greater habitat availability in northern areas.

In summary, this study challenges the assumption of a strong genetic isolation between populations of gypsum-associated plant species resulting from their island-like range. Indeed, long-term availability of suitable habitats appears to have permitted the preservation of high genetic variation and an active genetic interchange between populations of *N. hispanica*, at least based on plastid DNA markers. Future studies with determine the degree to which this pattern is also supported by genome-wide markers of this species, and by other gypsum-associated species from the western Mediterranean.

## Acknowledgements

This work would not have been possible without the support of Julián García, Rubén de Pablo, Jesús del Río, Leonardo Gutiérrez, Javier Puente, Javier Pavón and Luis M. Medrano, who helped us with field sample collection; Leopoldo Medina and Cyrille Chatelain, who granted us permission to use specimens from the MA and G herbaria respectively; and Emilio Cano, who provided his invaluable help in the RJB-CSIC molecular systematics laboratory.

## Supporting Information

**Table S1.** List of variables obtained to perform the niche modelling. Layers marked with an asterisk (*) were the ones selected to be used after a correlation test.

| Type | Layer | Type | Description | Source |
| --- | --- | --- | --- | --- |
| **Climatic information** | bio01 | Continuous | Annual mean temperature | CHELSA 1.2 |
|  | bio02 | Continuous | Mean diurnal range | CHELSA 1.2 |
|  | bio03 | Continuous | Isothermality | CHELSA 1.2 |
|  | bio04* | Continuous | Temperature seasonality | CHELSA 1.2 |
|  | bio05 | Continuous | Maximum temperature of warmest month | CHELSA 1.2 |
|  | bio06 | Continuous | Minimum temperature of coldest month | CHELSA 1.2 |
|  | bio07 | Continuous | Temperature annual range | CHELSA 1.2 |
|  | bio08* | Continuous | Mean temperature of wettest quarter | CHELSA 1.2 |
|  | bio09* | Continuous | Mean temperature of driest quarter | CHELSA 1.2 |
|  | bio10 | Continuous | Mean temperature of warmest quarter | CHELSA 1.2 |
|  | bio11* | Continuous | Mean temperature of coldest quarter | CHELSA 1.2 |
|  | bio12 | Continuous | Annual precipitation | CHELSA 1.2 |
|  | bio13 | Continuous | Precipitation of wettest month | CHELSA 1.2 |
|  | bio14 | Continuous | Precipitation of driest month | CHELSA 1.2 |
|  | bio15 | Continuous | Precipitation seasonality | CHELSA 1.2 |
|  | bio16* | Continuous | Precipitation of wettest quarter | CHELSA 1.2 |
|  | bio17* | Continuous | Precipitation of driest quarter | CHELSA 1.2 |
|  | bio18 | Continuous | Precipitation of warmest quarter | CHELSA 1.2 |
|  | bio19 | Continuous | Precipitation of coldest quarter | CHELSA 1.2 |
| **Edaphic information** | eda01 | Continuous | Bulk density (fine earth) | SoilGrids 2017 |
|  | eda02* | Continuous | Volumetric percentage of coarse fragments (>2 mm) | SoilGrids 2017 |
|  | eda03* | Continuous | Weight percentage of clay particles (<0.0002 mm) | SoilGrids 2017 |
|  | eda04 | Continuous | Weight percentage of silt particles (0.0002–0.05 mm) | SoilGrids 2017 |
|  | eda05* | Continuous | Weight percentage of sand particles (0.05–2 mm) | SoilGrids 2017 |
|  | eda06* | Continuous | Cation exchange capacity of soil | SoilGrids 2017 |
|  | eda07* | Continuous | Soil organic carbon content | SoilGrids 2017 |
|  | eda08 | Continuous | Soil organic carbon stock | SoilGrids 2017 |
|  | eda09* | Continuous | pH index measured in water solution | SoilGrids 2017 |
|  | eda10 | Continuous | pH index measured in KCl solution | SoilGrids 2017 |
| **Lithologic information** | iberlit* | Categorical | Cat. 01: Siliceous (igneous)<br>Cat. 02: Siliceous (metamorphic)<br>Cat. 03: Mafic-intermediate<br>Cat. 04: Ultramafic<br>Cat. 05: Clastic (coarse-grained)<br>Cat. 06: Clastic (fine-grained)<br>Cat. 07: Calcareous<br>Cat. 08: Dolomitic<br>Cat. 09: Gypsic<br>Cat. 10: Water | Fernández-Mazuecos & Glover (in prep.), based on Rodríguez Fernández et al. (2015) |

**Table S2.**
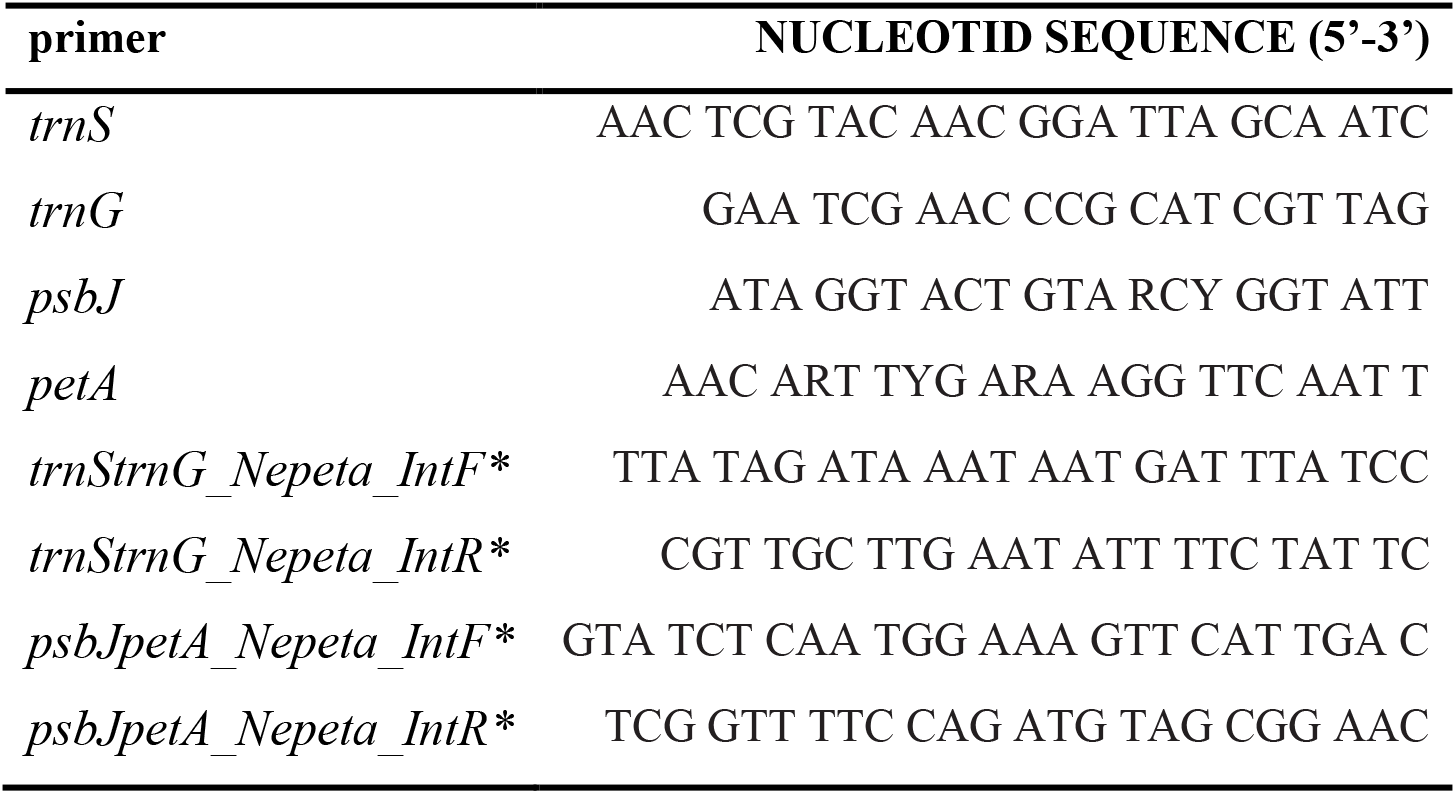
Nucleotide sequences of the primers used. Internal primers created *ad hoc* are marked with an asterisk.

**Table S3.** Origin of the genetic information for the samples used.

| Species (sample) | Origin | Analysis | GenBank<br>accession id |
| --- | --- | --- | --- |
| *Lallemantia peltata* | GenBank | Phylogeny | AJ420997 |
| *Dracocephalum kotschyi* | GenBank | Phylogeny | AJ420998 |
| *Dracocephalum grandiflorum* | GenBank | Phylogeny | AJ420999 |
| *Agastache pallida* | GenBank | Phylogeny | AJ421001 |
| *Agastache cana* | GenBank | Phylogeny | AJ421000 |
| *Nepeta wettsteinii* | GenBank | Phylogeny | AJ421038 |
| *Nepeta scrophularioides* | GenBank | Phylogeny | AJ515319 |
| *Nepeta kurdica* | GenBank | Phylogeny | AJ515320 |
| *Nepeta mussinii* | GenBank | Phylogeny | AJ515305 |
| *Nepeta congesta* var. *cryptantha* | GenBank | Phylogeny | AJ515161 |
| *Nepeta schiraziana* | GenBank | Phylogeny | AJ421034 |
| *Nepeta cataria* | GenBank | Phylogeny | AJ515313 |
| *Nepeta heliotropifolia* | GenBank | Phylogeny | AJ515312 |
| *Nepeta meyeri* | GenBank | Phylogeny | AJ421042 |
| *Nepeta saccharata* | GenBank | Phylogeny | AJ515314 |
| *Nepeta isaurica* | GenBank | Phylogeny | AJ515306 |
| *Nepeta fissa* | GenBank | Phylogeny | AJ421035 |
| *Nepeta balouchistanica* | GenBank | Phylogeny | AJ515606 |
| *Nepeta glomerulosa* | GenBank | Phylogeny | AJ515317 |
| *Nepeta mirzayanii* | GenBank | Phylogeny | AJ515309 |
| *Nepeta straussii* | GenBank | Phylogeny | AJ421040 |
| *Nepeta gloeocephala* | GenBank | Phylogeny | AJ515308 |
| *Nepeta bornmuelleri* | GenBank | Phylogeny | AJ515310 |
| *Nepeta hormozganica* | GenBank | Phylogeny | AJ515160 |
| *Nepeta denudata* | GenBank | Phylogeny | AJ515304 |
| *Nepeta ispahanica* | GenBank | Phylogeny | AJ515318 |
| *Nepeta pungens* | GenBank | Phylogeny | AJ421036 |
| *Nepeta eremophila* | GenBank | Phylogeny | AJ515315 |
| *Nepeta cephalotes* | GenBank | Phylogeny | AJ421037 |
| *Nepeta crassifolia* | GenBank | Phylogeny | AJ515307 |
| *Nepeta sibirica* | GenBank | Phylogeny | AJ421039 |
| *Nepeta pogonosperma* | GenBank | Phylogeny | AJ421041 |
| *Nepeta menthoides* | GenBank | Phylogeny | AJ421002 |
| *Nepeta crispa* | GenBank | Phylogeny | AJ515159 |
| *Nepeta binaloudensis* | GenBank | Phylogeny | AJ515311 |
| *Nepeta assurgens* | GenBank | Phylogeny | AJ515316 |
| *Nepeta oxyodonta* | GenBank | Phylogeny | AJ420996 |
| *Nepeta laxiflora* | GenBank | Phylogeny | AJ420995 |
| *Nepeta hispanica* (MA855920) | MA Herbarium | Phylogeny | Forthcoming |
| *Nepeta hispanica* (MA907269) | MA Herbarium | Phylogeny | Forthcoming |
| *Nepeta ucranica* (MA744790) | MA Herbarium | Phylogeny | Forthcoming |

**Table S4.**
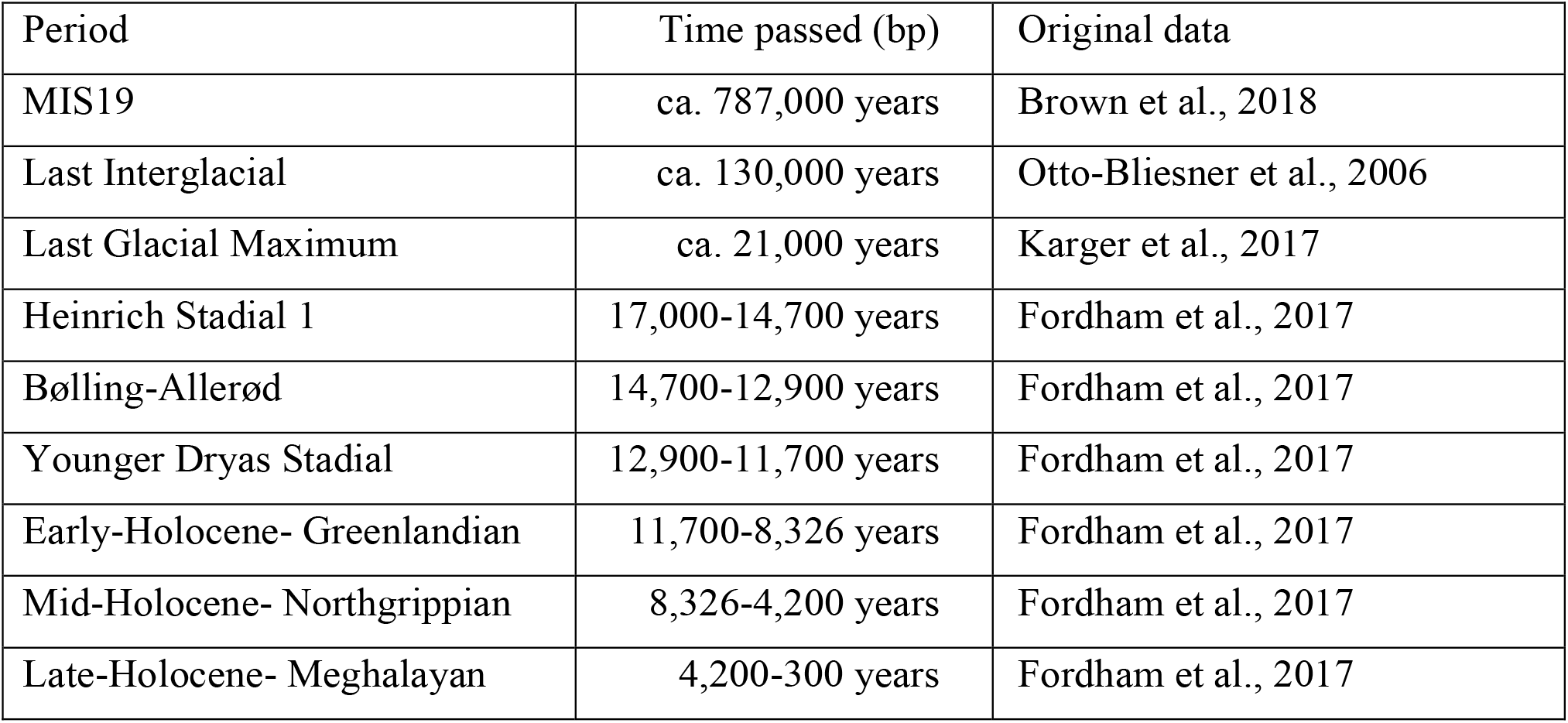
Periods projected to the past to evaluate environmental suitability.

